# NPC1L1 inhibition disturbs lipid trafficking and induces large lipid droplet formation in intestinal absorptive epithelial cells

**DOI:** 10.1101/2020.04.17.036814

**Authors:** Takanari Nakano, Ikuo Inoue, Yasuhiro Takenaka, Rina Ito, Norihiro Kotani, Sawako Sato, Yuka Nakano, Masataka Hirasaki, Akira Shimada, Takayuki Murakoshi

**Author notes:** To whom correspondence should be addressed: Takanari Nakano, Ph.D., Department of Biochemistry, Faculty of Medicine, Saitama Medical University, 38 Morohongo, Moroyama, Iruma-gun, Saitama 350-0495, Japan.

## Abstract

Ezetimibe inhibits Niemann-Pick C1-like 1 (NPC1L1) protein, which mediates intracellular cholesterol trafficking from the brush border membrane to the endoplasmic reticulum, where chylomicron assembly takes place in enterocytes or in the intestinal absorptive epithelial cells. Cholesterol is a minor lipid component of chylomicrons; however, whether or not a shortage of cholesterol attenuates chylomicron assembly is unknown. The aim of this study was to examine the effect of NPC1L1 inhibition on trans-epithelial lipid transport, and chylomicron assembly and secretion in enterocytes. Caco-2 cells, an absorptive epithelial model, grown onto culture inserts were given lipid micelles from the apical side, and chylomicron-like triacylglycerol-rich lipoprotein secreted basolaterally were analyzed after a 24-h incubation period in the presence of ezetimibe up to 50 μM. The secretion of lipoprotein and apolipoprotein B48 were reduced by adding ezetimibe (30%, *p*<0.01 and 34%, *p*<0.05, respectively). Additionally, ezetimibe accelerated intracellular apoB protein degradation by approximately 2.8-fold and activated sterol regulatory element binding protein 2 by approximately 1.5-fold: These are indicators whether the cells are sensing cellular cholesterol shortage. Thus, ezetimibe appeared to limit cellular cholesterol mobilization required for lipoprotein assembly. In such conditions, large lipid droplet formation in Caco-2 cells and the enterocytes in mice were induced, implying that unprocessed triglyceride was sheltered in these compartments. Although ezetimibe did not reduce the post-prandial lipid surge appreciably in triolein-infused mice, the results of the present study indicated that NPC1L1-mediated supply chylomicron with cholesterol may participate in a novel regulatory mechanism for the efficient chylomicron assembly and secretion.

## INTRODUCTION

The small intestine secretes chylomicron, or a triacylglycerol (TAG)-rich lipoprotein, to deliver alimentary fatty acids into the circulation [1]. Lipid in chylomicron is mainly composed of TAG together with the other minor lipids; phospholipids and cholesterol. Limitations in the supply of the minor lipids can therefore disturb TAG delivery. Biliary phospholipid supply to the lumen is required for efficient chylomicron secretion [2]. Such effects were also observed in *ABCB4*-knockout mice that lack bile-mediated phospholipid supply to the lumen [3]. Moreover, endogenous phospholipid synthesis in the intestinal epithelia was needed when a high-fat diet was fed to mice [4].

Cholesterol supply may also affect chylomicron secretion efficiency [5]. Mice treated with statins showed a greater post-prandial TAG surge in the early phase after a bolus of lipid in rodents (0–3 h) [6, 7], in which conditions it was indicated that cholesterol synthesis was increased in the small intestine [8]. Similarly, such a post-prandial TAG surge was observed in mice with greater intestinal cholesterol absorption efficiency in mice disrupted for cholesterol efflux transporters, ATP-binding cassette G5/G8 [9]. Moreover, activation of liver X receptor attenuated cholesterol absorption, stimulated lipid droplet formation, and retarded TAG transport in the intestine of Zebra fish [10, 11]. On the other hand, cholesterol depletion by statins or an acyl-CoA acyltransferase (ACAT) inhibitor in lipoprotein-producing cell lines, Caco-2 and HepG2 cells, accelerated apolipoprotein (apo) B degradation [12, 13]. These observations suggested that cholesterol supply is involved in the regulation of chylomicron assembly and secretion.

The brush border membrane (BBM) of enterocytes is rich in cholesterol [14]. Field et al. [15] showed that plasma membrane cholesterol is the major source for chylomicron-like TAG-rich lipoprotein in Caco-2 cells. Niemann-Pick C1-like 1 (NPC1L1) mediates cholesterol trafficking from the BBM to the endoplasmic reticulum (ER) [16–18]. Intestinal *NPC1L1* gene expression is positively correlated with cholesterol content in circulating chylomicrons [19], suggesting that NPC1L1-mediated cholesterol supply to chylomicron formation plays a role in the lipid assimilation process in enterocytes. Studies with experimental animals supported this possibility: In mice fed a Western diet, ezetimibe ameliorated excess lipoprotein secretion in the small intestine [20]; and in hamsters ezetimibe treatment decreased serum apoB48 protein levels in a post-prandial state even when the animals were treated with Tyloxapol, a lipoprotein lipase inhibitor used to block chylomicron clearance from the circulation [21]. With this treatment, extra-intestinal effects for clearance were minimized.

In the present study, we examined the effect of ezetimibe, a prescribed drug that inhibits NPC1L1 for the treatment of hypercholesterolemia, on lipid transport efficiency and cytosolic lipid droplet formation to reveal the direct effect of NPC1L1 inhibition on epithelial lipid mobilization in enterocyte-like differentiated Caco-2 cells and the proximal small intestine of C57BL mice.

## MATERIALS AND METHODS

### Reagents

Pluronic L81® (BASF Co., Washington, NJ, USA), a hydrophobic surfactant that inhibits chylomicron secretion, was a kind gift from Prof. Patrick Tso (Dept. of Pathology and Laboratory Medicine, University of Cincinnati, OH, USA). 9, 10(n)-^3^H-oleic acid (specific activity, 45.5 Ci/mmol) and [4-^14^C]-cholesterol (specific activity, 49.8 mCi/mmol) were purchased from GE Healthcare (Piscataway, NJ, USA). 1-^14^C-oleic acid was obtained from Moravek Biochemicals Inc. (cat.no. MC 406, 55 mCi/mmol, Brea, CA, USA). Other reagent information was shown in Table S1.

### Antibodies

The antibodies used in the present study are listed in Table S2.

### Differentiation of Caco-2 cells on filter membranes

Caco-2 cells (a gift from Dr. Sylvie Demignot, INSERM, Paris, France) were differentiated on filter membranes (Individual Cell Culture Inserts with 0.4 μm pore size, BD Biosciences, San Jose, CA, USA) [22, 23]. Briefly, Caco-2 cells were plated at a density of 5 × 10^4^ cells per cm^2^ membrane and grown in a humidified atmosphere containing 5% CO_2_, at 37°C in Dulbecco’s modified Eagle’s medium (DMEM) with 4.5 g/L glucose and GlutaMAX™ (Product No. 35050061, ThermoFisher Scientific, Carlsbad, CA, USA) or 20% heat-inactivated fetal calf serum (FCS). After 1 week, the cells were cultured in asymmetric conditions, with DMEM containing 1.0 g/L glucose and GlutaMAX™ in the upper compartment (apical media) and DMEM containing 1.0 g/L glucose and 20% FCS in the lower compartment (basolateral media) for 1 week. Then, the basolateral medium was replaced with medium containing ITS supplement (Life Technologies) and cultured for an additional week.

### Preparation of lipid micelles

Lipid micelles were prepared as described previously [22, 23] with some modifications. In brief, stock solutions of oleic acid, lysophosphatidylcholine, and non-esterified cholesterol were mixed in a sterile glass tube and dried. The resulting dried lipids were dissolved in 83 μl of a sterile solution of 24 mM taurocholate. Then 1 ml of serum-free medium was added. The final lipid concentrations were 0.6 mM oleic acid, 0.05 mM cholesterol, 0.2 mM lysophosphatidylcholine, and 2 mM taurocholate, which mimics post-digestive duodenal micelles [24]. Lipid micelles were used within 2 h after preparation.

### Lipoprotein secretion by differentiated Caco-2 cells

Two hours before the addition of lipid micelles, differentiated Caco-2 cell monolayers were treated with ezetimibe (0.5–50 μM) or vehicle (1% v/v ethanol). We employed these dosages of ezetimibe because earlier studies used submillimolar dosages (20–150 μM) [16, 25, 26]. Freshly prepared lipid micelles (1.5 ml) were added to the upper compartment of a culture insert with or without ezetimibe. The basolateral media was also changed to ITS-containing FCS-free media. Then, the cells were incubated for 24 h. For tracer experiments, ^3^H-oleic acid and ^14^C-cholesterol were added with final radioactivity levels of 2 μCi/ml and 1 μCi/ml, respectively. After incubation with the tracer-containing lipid micelles, the media were harvested, and the cells were rinsed twice with ice-cold phosphate-buffered saline (PBS), scraped into 0.5 ml of a lysis buffer (1% Triton X-100, 5 mM EDTA in PBS) supplemented with 2% protease inhibitor cocktail, disrupted using a 23G needle and syringe, and immediately frozen at –80°C until analysis.

Lipoproteins secreted into the basolateral media were separated from free tracers using a PD-10 desalting column (GE Healthcare) as described previously [23]. Fifty μl of each elution fraction was suspended in 1 ml of liquid scintillation reagent (Ultima Gold, PerkinElmer, Wellesley, MA, USA), and the radioactivity in each vial was measured using a scintillation counter. The distribution of radioactivity (%) was calculated by comparing the sum of whole counts in the media and cell lysates (total count).

### Apolipoprotein and lipid assays

The concentration of apoB48 was determined using an apoB48 CLEIA Fujirebio kit and a LUMIPULSE system (Fujirebio, Tokyo, Japan) [27]. Total cholesterol (both non-esterified and esterified) and TAG concentrations were determined using a Cholesterol/Cholesteryl Ester Quantitation Kit and a Triglyceride Quantification Kit (Biovision Inc. San Francisco, CA. USA), respectively.

### Western blotting

Cell lysates were separated on 5% and 8% polyacrylamide gels for apoB and SREBPs, respectively, and transferred to polyvinylidene fluoride membranes (Amersham Biosciences). For the antibodies used, see Table S2. Respective bands were visualized by chemiluminescence (ECL™, Western Blotting Detection Reagents; BD Biosciences) using a high-performance chemiluminescence film (Hyperfilm™ ECL™, BD Biosciences). The films were scanned, and the density of each band was translated into arbitrary units using ImageJ software (http://rsb.info.nih.gov/ij/).

### Metabolic labeling of Caco-2 cells and immunoprecipitation of apolipoprotein B

Differentiated Caco-2 cells were used for pulse-chase experiments as described previously [28, 29] with some modifications. The cells were treated with or without 50 μM ezetimibe for 2 h, lipid micelles were added to the apical media, and the cells were incubated for 2 h. Then, the cells were depleted of methionine and cysteine for 30 min by placing in methionine- and cysteine-free medium (methionine cysteine free DMEM, Cat. No. 21013-024, Life Technologies) with or without 50 μM ezetimibe. Then, newly synthesized proteins in the cells were labeled by treating with 150 μCi/ml of ^35^S-methionine/cysteine (EXPRE^35^S^35^S Protein Labeling Mix, NEG772, PerkinElmer) apically for 45 min (pulse labeling). Complete protein synthesis of the cells was allowed by replacing the apical and basolateral media with methionine- and cysteine-containing 0.75 ml of fresh ITS-containing FCS-free media, which started the chase phase for 20 min or 2 h.

ApoB proteins in the cell lysate or basolateral culture supernatants were immunoprecipitated using anti-apoB antibodies (Table S2) and IP kit-Dynabeads Protein G (Veritas, Tokyo, Japan). Linear (5%) or gradient 5–15% gels were used to separate precipitated apoB by SDS-PAGE, and dried. The gels were exposed to imaging plates and the images of separated tracer-labeled proteins were obtained by reading the plate using a Typhoon 9410 (GE Healthcare) image analyzer. The density of the bands was determined using ImageJ. We estimated apoB protein secretion from the resulting density.

### Quantitative RT-PCR

We obtained total RNA samples from differentiated Caco-2 cells exposed to ezetimibe (50 μM) for 6 or 24 h without lipid micelles or FCS using ISOGEN (Wako Pure Chemical Industries, Tokyo, Japan). These RNA samples were reverse-transcribed using a QuantiTect Rev. Transcription Kit (QIAGEN, Venlo, Netherlands). Quantitative expression analysis was performed using an ABI PRISM 7900HT Sequence Detector (Life Technologies) with SYBR green technology. The cycle threshold (Ct), corresponding to the number of cycles after which the target-DNA concentration increase becomes exponential, was monitored. Results were analyzed using SDS 2.1 Software (Life Technologies). All reactions were performed in duplicate for each sample. We used *HPRT1* as a housekeeping gene. The primer sets used are listed in Table S3.

### Lipid droplet staining for Caco-2 cells

Lipid droplets in cells were visualized with BODIPY™ 493/503. Caco-2 cell monolayers were fixed with 4% paraformaldehyde (PFA) for 10 min at room temperature, stained with BODIPY (1 μM) for 15 min, washed again, and mounted with PBS containing DAPI (5 μg/ml). Fluorescence images were obtained immediately using a confocal microscope (LSM 710, Carl Zeiss, Oberkochen, Germany). Z-stack images were obtained at 1-μm intervals.

### Mice

Eight-weeks-old C57BL/6J male mice were purchased from Clea Co., Ltd. (Tokyo, Japan) and used at 10–12-weeks of age. They were given standard chow and housed in cages in a temperature-controlled room with 12-h light cycling. All animal experiments were approved by the Animal Care Committee of Saitama Medical University.

### Preparation of luminal perfusate

Krebs-HEPES buffer containing gall (2% w/v) and pancreatin (2% w/v) was vigorously mixed with oleic acid (20% v/v). The aqueous layer was recovered as perfusate as below. The concentration of non-esterified fatty acid in the perfusate was measured using LabAssay™ NEFA (Wako). ^14^C-oleic acid was added to a final concentration of 1 μCi/ml when necessary.

### Intestinal lipid perfusion

Luminal perfusion assays were performed in accordance with van der Velde et al. [30] with some modifications [31]. Ezetimibe was given orally to C57BL/6J mice at indicated doses the day before the assay (during from –16 to –18 h for perfusion), because ezetimibe is glucuronidated in the intestine and liver to exert the pharmacological activity in the BBM of the small intestine as well as the parent compound. Ezetimibe was given again 3 h before the assay. Dosage of ezetimibe was shown as the total below: The sum of the twice administration. The jejunum was exposed by midline incision, externalized, and prepared for perfusion by cannulating at the inlet and the outlet of the upper small intestinal segment under anesthesia with a cocktail of ketamine (65.2 mg/kg) and xylazine (18.5 mg/kg). After flushing the luminal content using Krebs-HEPES buffer (0.25 ml/min) with an infusion pump (Harvard Apparatus Inc., Holliston, MA, USA), luminal perfusate was infused for 30 min (0.05 ml/min). After the lumen and the blood in the circulation were flushed out with saline, the perfused small intestinal segments were stored for tracer counting [31], or fixed with 4% PFA for microscopic analysis.

### Fluorescence microscopic analysis

PFA-fixed intestinal segments were submerged into sucrose-containing PBS (serially to 10%, 15%, 20% w/v sucrose), and snap-frozen with OCT compound® (Sakura® Finetek Japan, Tokyo, Japan). Four-μm cryosections were stained with BODIPY™ 493/503 and DAPI as described above. ApoB was immunostained with the antibodies listed in Table S2. DAPI and Alexa Fluor™ 594 Phalloidin were used for counterstaining.

### Transmission electron microscopy

After intestinal lipid perfusion for 30 min, the perfused segments were fixed with 2.5% glutaraldehyde for 1 day and post-fixed in a 1% OsO_4_ solution buffered with Sorenson’s phosphate at pH 7.4. Then, the segments were rapidly dehydrated in alcohol and embedded in Epon 812. Thin sections of representative areas were then examined using an electron microscope (JEM-1400 Transmission Electron Microscope, JEOL Ltd., Tokyo, Japan).

### Triolein-induced TAG surge in mice

We gave 50 μg ezetimibe in 100 μl of 10% DMSO orally to 10–12-weeks-old C57BL/6J male mice and fasted for 3 h (10 AM to 1 PM). Then we gave 0.2 ml triolein containing 50 μg ezetimibe in 100 μl of 10% DMSO again and Tyloxapol (500 μg/kg), a lipoprotein lipase inhibitor, via intraperitoneal injection. Thus, the total dose of ezetimibe was 100 μg per mouse. Three hours after the injections, blood was collected for TAG assays. We gave BLT-1 (1 mg) and orlistat (2 mg) with triolein to mice orally as well.

### Statistical analysis

Data are shown as means ± SEM of four replicate assays unless indicated otherwise. Means were compared using Student’s *t*-test or the Mann–Whitney *U* test if the data were distributed non-parametrically. For multiple comparisons, we used Dunnett’s test. Statistical analyses were performed using statistical software (JMP® 13.2.1, SAS Institute).

## RESULTS

### Ezetimibe reduces chylomicron secretion in differentiated Caco-2 cells

At first, we validated differentiated Caco-2 cells for TAG-rich lipoprotein secretion upon lipid exposure to Pluronic L81®, a potent chylomicron synthesis inhibitor [32] (Fig. 1A). Tracing ^3^H-oleic acid in the culture showed that treatment with Pluronic L81 abolished lipid micelle-induced TAG-rich lipoprotein secretion (Fig. 1B), confirming that the cells utilize chylomicron assembly pathways for the secretion. The addition of ezetimibe reduced secretion in a dose-dependent manner by more than 30% at a 50 μM dosage. The reduction was also confirmed by an HPLC analysis (Fig. S1). On the other hand, ezetimibe treatment did not show appreciable cytotoxic features (Fig. S2),

**Figure 1.**
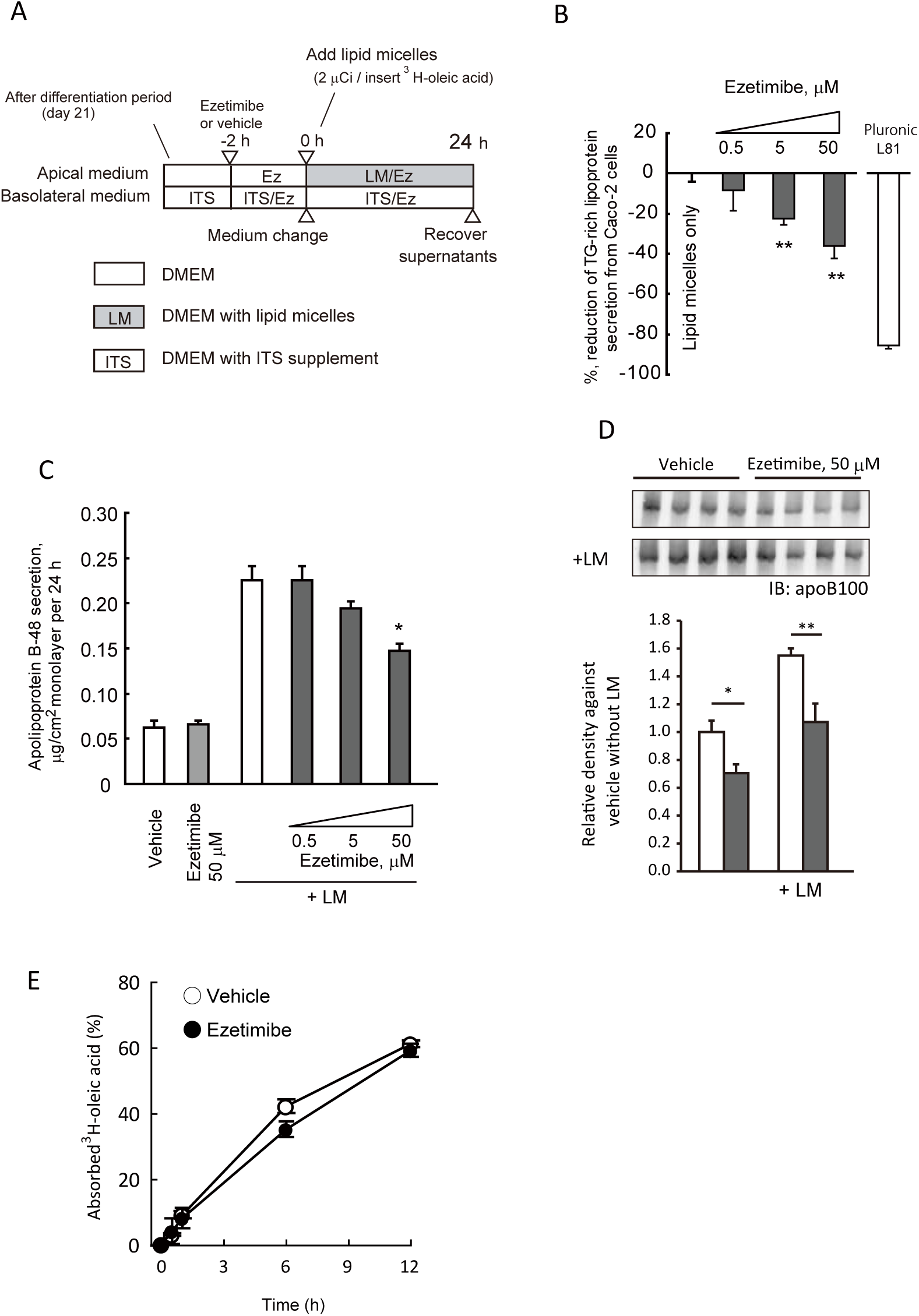
Ezetimibe inhibits lipoprotein secretion in differentiated Caco-2 cells. *A*, Schematics of the protocol for lipoprotein secretion by differentiated Caco-2 cells. Hours (h) indicate relative time periods to that when lipid micelles were added. Ez, ezetimibe; ITS, supplemented with insulin, transferrin, and selenium instead of fetal calf serum; LM, lipid micelles. *B*, Secretion of newly synthesized lipoprotein into the basolateral media was determined by tracing the radioactivity of ^3^H-oleic acid. Pluronic L81 (10 μg/ml) was used as an inhibitor of chylomicron secretion. ** *p* < 0.01 vs. control with lipid micelles only by Dunnett’s test. Data are shown as mean and SEM of four replicate assays. *C and D*, Ezetimibe reduced secretion of apolipoprotein (apo) B. *C*, Ezetimibe reduced apoB48 secretion by Caco-2 cells in the presence of lipid micelles. ApoB48 protein levels were determined using an ELISA. * *p* < 0.05 vs. lipid micelles only by Dunnett’s test. +LM indicates the addition of lipid micelles to the apical media. Data are shown as mean and SEM of four replicate assays. *D*, Ezetimibe reduced apoB100 secretion by Caco-2 cells, as determined by Western blotting analysis. *, *p* < 0.05; **, *p* < 0.01 vs. vehicle by Student’s *t*-test. Data are mean and SEM of four replicate assays. *Open bars*, vehicle; *gray bars*, ezetimibe (50 μM) *E*, Ezetimibe had little effect on medium-to-cell ^3^H-oleic acid transit in differentiated Caco-2 cells. *Open circles*, vehicle (1% ethanol); *closed circles*, 50 μM ezetimibe. Data are shown as mean ± SEM of triplicate assays.

### Ezetimibe attenuates apoB protein secretion

Upon lipid micelle exposure, the structural protein apoB48 of chylomicrons in the basolateral culture supernatants increased approximately 4-fold; ezetimibe attenuated the increase by 34% (Fig. 1C). In the absence of lipid micelles, the treatment did not alter the basal apoB48 secretion. For the very low density lipoprotein structural protein apoB100, the secretion was increased 1.5-fold with lipid micelles; the increase was attenuated by 31% by ezetimibe. ApoB100 secretion was also reduced by 29% in the absence of lipid micelles (Fig. 1D); unlike the case of apoB48 (Fig. 1C). No significant difference was observed in mRNA abundance with ezetimibe treatment (Fig. S3). Ezetimibe did not affect oleic acid and cholesterol uptake by the cells [Fig. 1E for oleic acid; data in Ref. [31] for cholesterol].

### Intracellular apolipoprotein B48 degradation was accelerated in the presence of ezetimibe

Disruption of cellular cholesterol metabolism in Caco-2 cells [29, 33] or HepG2 cells [12, 13] accelerated intracellular apoB protein degradation. To estimate the degradation in the presence of ezetimibe, we performed pulse-chase analysis followed by apoB immunoprecipitation (Fig 2A).

**Figure 2.**
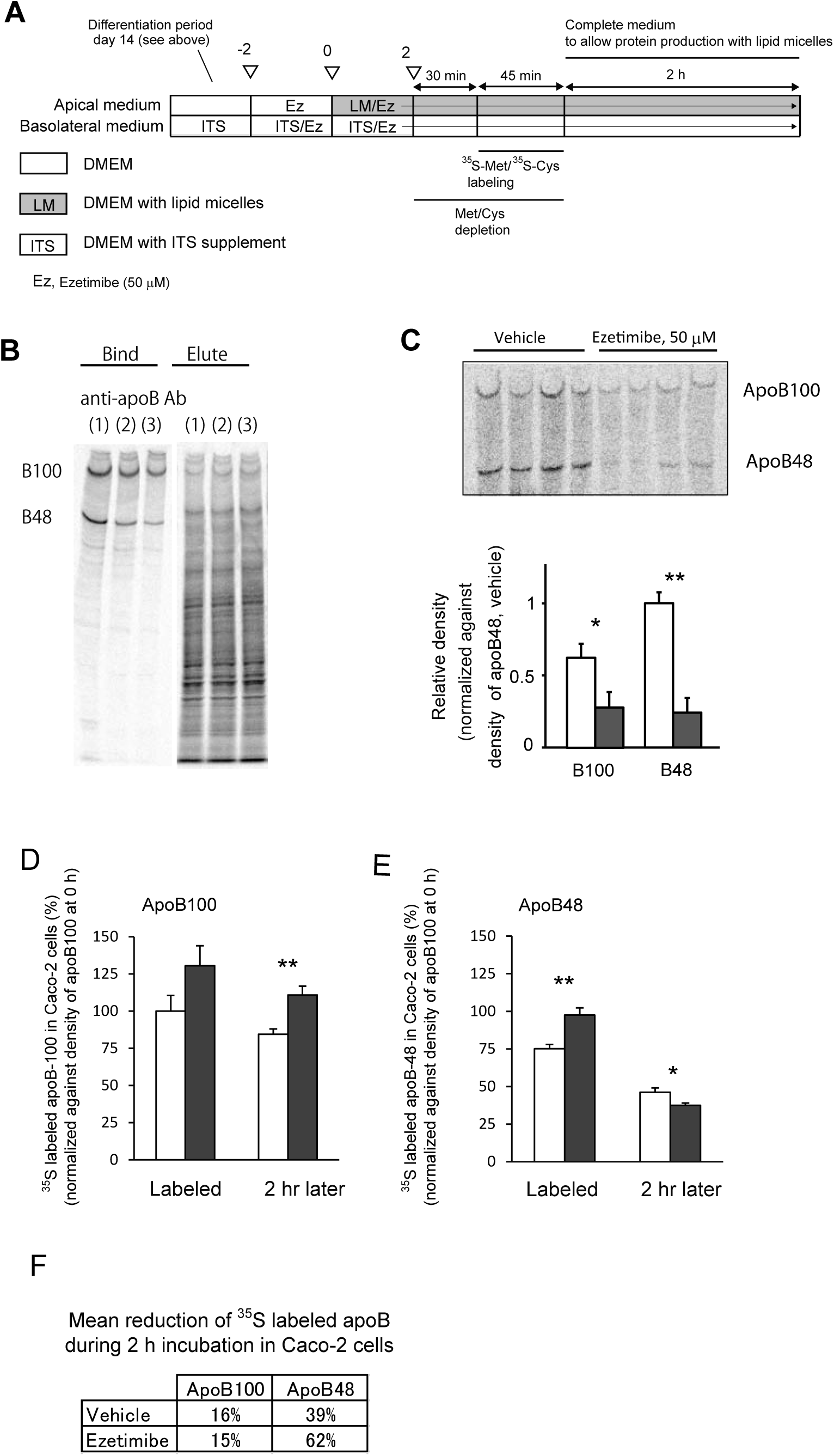
*A*, Schematics of the protocol for pulse-chase labelling for newly synthesized apolipoprotein (apo) B. *B*, ApoB-immunoprecipitation for Caco-2 cell lysate. Numbers in parentheses indicate anti-apoB antibody used (see *Materials and Methods*). The left one (1) showed better recovery and specificity for apoB proteins among the three and was used for further experiments. Proteins were separated using a 5–15% polyacrylamide gradient gel. *C*, Pulse-chase analysis showed that ezetimibe attenuated newly synthesized apoB48 and B100 secretion. The upper panel shows density patterns of secreted apoB. Data are shown as mean and SEM of four replicate assays. *, *p* < 0.05; **, *p* < 0.01 vs. vehicle by Student’s *t*-test. Proteins were separated using a 5% polyacrylamide gel. *D and E*, Ezetimibe increased apoB protein synthesis and accelerated the degradation. Newly synthesized apoB100 (*D*) and apoB48 (*E*) are increased approximately 2- and 1.5-fold, respectively, during the 0–20 min period after the pulse. *F*, Percent reductions during from 20 min to 2 h after the pulse (data from *D* and *E*) were calculated and shown. The degradation of apoB100 and apoB48 proteins were approximately 1.3- and 2.8-fold, respectively, with ezetimibe.

First, we selected anti-apoB antibody #1, which showed higher recovery of apoB protein by immunoprecipitation among the three antibodies tested (Fig. 2B). When we analyzed the basolateral supernatants obtained after a 2-h incubation period, ezetimibe attenuated newly synthesized apoB100 and apoB48 secretion by 57% and 77%, respectively (Fig. 2C). These reductions were approximately twofold greater than that for total apoB secretion (Fig. 1). When we estimated apoB protein production rate at 20 min of the chase phase, apoB48 and B100 protein synthesis were increased in the presence of ezetimibe (Figs. 2D-E). Loss of labeled apoB proteins in the cells at 20 min and 2 h after the pulse were approximately 2.8-fold for apoB48 (Fig. 2E; 16.5% vs. 6.7%, see the inset *F* in Fig. 2) and approximately 1.3-fold for apoB100 (Fig. 2D; 6.3% vs. 4.0%) with ezetimibe, showing a greater effect of ezetimibe on apoB48 degradation.

### Ezetimibe activates SREBP2 in differentiated Caco-2 cells

We showed previously that ezetimibe does not limit medium-to-cell cholesterol transit in differentiated Caco-2 cells [31]. On the other hand, consistent with two earlier reports [15, 34], we observed that cholesterol in TAG-rich lipoproteins secreted from Caco-2 cells was of endogenous origin (Fig. S4). Collectively, ezetimibe can affect intracellular cholesterol metabolism or trafficking. Indeed, ezetimibe reduced cholesterol esterification in the ER, in which the availability of cholesterol for cells is being monitored [16].

To estimate cellular response to ezetimibe treatment, we examined whether or not ezetimibe activates sterol regulatory element binding protein (SREBP) 2, a component of the sterol sensor in Caco-2 cells in the absence of lipid micelles (Fig. 3A). This protein is cleaved when there is a shortage of available cholesterol [35] and releases nuclear type SREBP2 to compensate the shortage by inducing genes related to *de novo* synthesis of cholesterol, for example. Western blotting analysis showed that ezetimibe activated the cleavage of SREBP2, increasing the nuclear type by 1.5-fold both at 6 h and 24 h (Figs. 3B-C), but not SREBP1, a regulator of fatty acid synthesis. Quantitative RT-PCR analysis showed that the expression levels of target genes of SREBP2 were modulated by ezetimibe treatment. Of the six genes examined, five showed corresponding changes (Fig. 3D). Activation of SREBP2 down-regulated *MTTP* gene expression, consistent with a previous report [36]. In contrast, only one (*FAS*) of the six genes targeted by SREBP1 showed an appreciable change (Fig. 3E).

**Figure 3.**
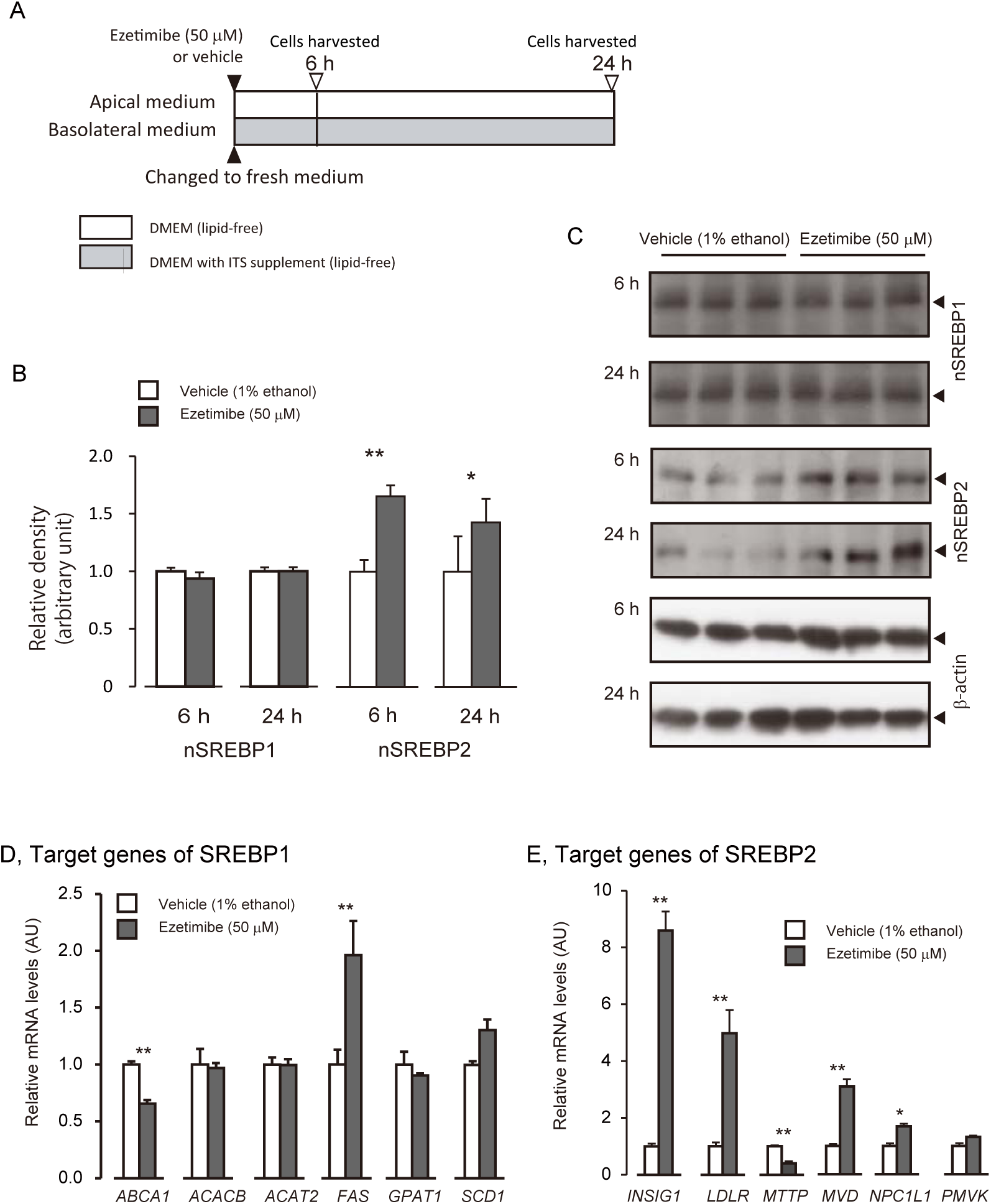
Ezetimibe increases the nuclear type of sterol regulatory element binding protein 2 (nSREBP2), the active form of the transcription factor. A, Schematics of the protocol for quantitative RT-PCR analysis. Hours (h) indicate time periods when the cells harvested. Ez, ezetimibe; ITS, supplemented with insulin, transferrin, and selenium instead of fetal calf serum. *B-C*, Western blotting analysis of cellular nSREBP1 and nSREBP2 after either 6 h or 24 h incubation with ezetimibe. Data are shown as the mean and SEM of triplicate assays. Comparison of the densities in *B*. *, *p* < 0.05; **, *p* < 0.01 (Student’s *t*-test). *C* shows the membrane images. *D* and *E*, Changes in mRNA abundance of SREBP1 (*D*) and SREBP2 (*E*) target genes. Of the six mRNA species tested, five changed significantly by adding ezetimibe in the SREBP2 target genes (*E*): the SREBP1 target genes did not expect for *FAS. ABCA1*, ATP-binding cassette A1; *ACACB*, Acetyl-CoA carboxylase β; *ACAT2*, Acetyl-CoA acetyltransferase 2; *FAS*, Fatty acid synthase; *GPAT1*, 1-acylglycerol-3-phosphate *O*-acyltransferase 1 (lysophosphatidic acid acyltransferase, αα); *SCD1*, Stearoyl-Coenzyme A desaturase 1; *INSIG1*, Insulin-induced gene 1 protein; *LDLR*, low density lipoprotein receptor; *MTTP*, microsomal triglyceride transfer protein; *MVD*, mevalonate decarboxylase; *PMVK*, phosphomevalonate kinase; *NPC1L1*, Niemann-Pick C1-like protein 1. * *p* < 0.05, **, *p* < 0.01 vs. corresponding vehicle control by Student’s *t*-test. Data shown are the mean and SEM of four replicate assays.

### Ezetimibe accelerates large lipid droplet formation in Caco-2 cells

Lipid droplets provide buffering spaces for chylomicron production [37–40]. They function in the storage of lipids before they are packed into chylomicrons. Because TAG-rich lipoprotein production was impaired in ezetimibe-treated Caco-2 cells (Fig. 1), we tried to examine whether ezetimibe affects lipid droplet formation (Fig. 4A). Fluorescent microscopy with BODIPY staining showed that ezetimibe induced large lipid droplet formation even in lipid-free conditions (Fig. 4B, *upper right panel*; Fig. S5A). This occurred even with lower doses of ezetimibe (Fig. S5B). Then we added lipid micelles and found that larger lipid droplets developed with time (Fig. 4B, *right panels*). With ezetimibe treatment, most lipid droplets were localized at the apical perinuclear regions (Figs. 4C-E). This effect was prominent with inhibitors for ER- and Golgi-associated trafficking (Fig. 4F; nocodazole and tunicamycin for ER-to-Golgi trafficking; colchicine for the secretory pathway; and monensin for the trans-Golgi function) and recapitulated weakly with CP-346086, which inhibits microsomal triglyceride transfer protein (MTP), a key enzyme for lipoprotein assembly. The other three inhibitors for lipid metabolism (HMG-CoA, acyl-coenzyme A:cholesterol acetyltransferase 2, or microsomal triglyceride transfer protein) did not show any comparable effect to ezetimibe (Fig. S5C).

**Figure 4.**
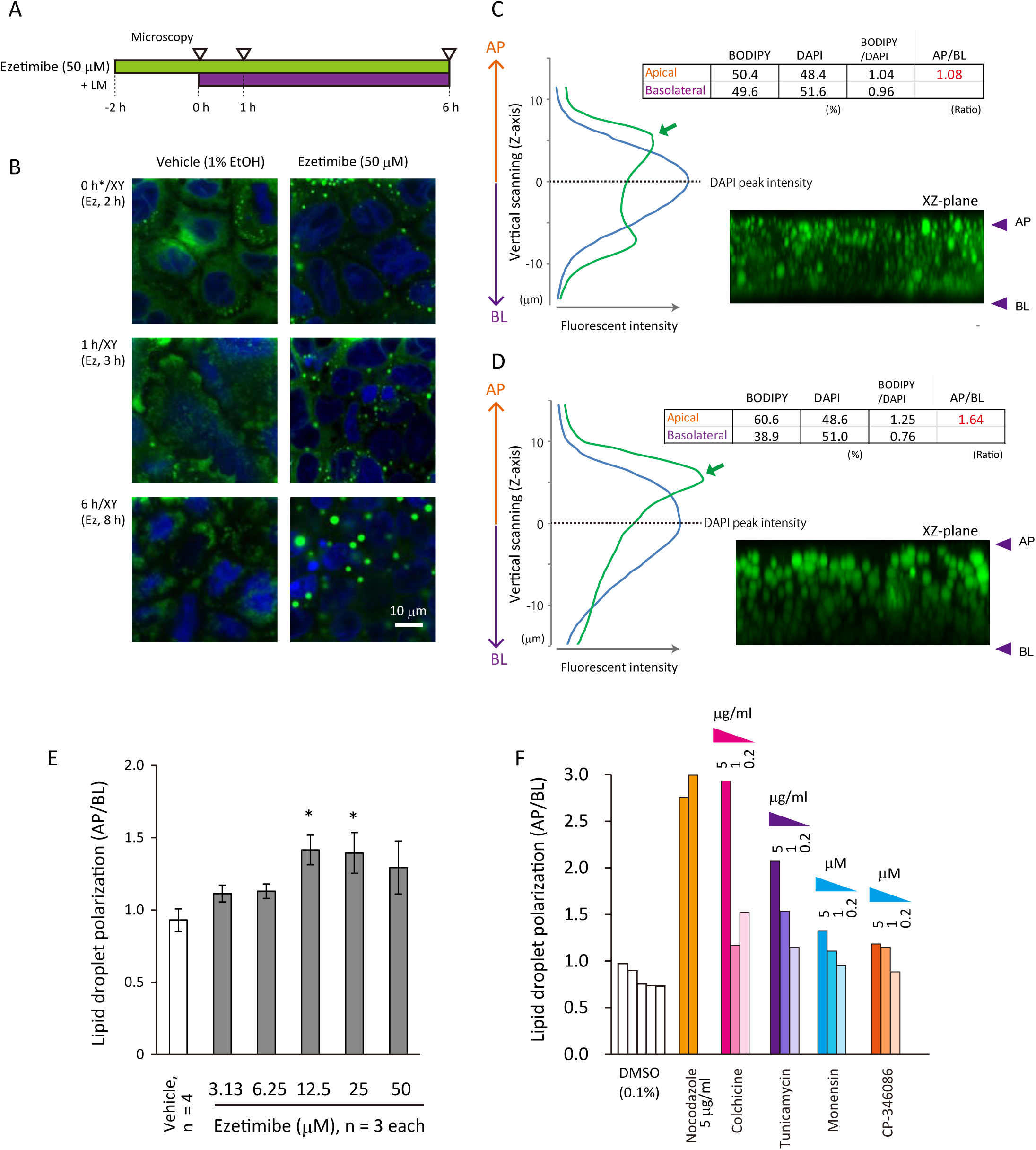
Ezetimibe induces large lipid droplet formation on the apical side of the nuclei in differentiated Caco-2 cells. *A* indicates the periods of chemical treatments and the microscopic analysis performed. The panels in *B* show the representative time-dependent enlargement of lipid droplets in ezetimibe-treated cells. *C* and *D*, Representatives of Z-axis confocal microscopy scanning of Caco-2 cells after 6-h lipid micelle exposure (the bottom in *B*). Green and blue lines show arbitrary fluorescent intensity of BODIPY and DAPI, respectively. The plane with the highest intensity of DAPI was set as 0 μm. Then, the upper and the lower were assumed as apical (AP) and basolateral (BL) sides, respectively. The two fluorescent signal counts were divided into the two areas and shown as percentage in the *Insets*. The fluorescent intensity of BODIPY was normalized by that of DAPI. Normalized values were used to estimate polarization of BODIPY-staining (AP/BL in *red*). XZ-plane confocal images for BODIPY were shown without the DAPI signals. With ezetimibe, BODIPY intensity was increased at the apical perinuclear region (*Green arrows*). *E*, Differentiated Caco-2 cells accumulate lipid droplets in the perinuclear region in the presence of ezetimibe. The values of AP/BL were calculated as shown in *C* and *D*. Data are shown as mean and SEM of four replicate (vehicle, 1% ethanol) or triplicate assays (ezetimibe-treated). *, p < 0.05 by the Dunnett’s test. *F*, ER- or Golgi-associated trafficking inhibitors induced apical lipid droplet accumulation in differentiated Caco-2 cells. Each bar indicates an individual experiment (n = 1). The Y-axis shows the same scale with *E.* CP-346086 is a microsomal triglyceride transfer protein inhibitor.

### Ezetimibe stimulates the formation of large lipid droplets and disturbs apoB localization in murine enterocytes

The prepared lipid containing perfusate (Fig. 5A) contained 15.5 ± 0.8 (mean ± standard deviation) mEq/L of non-esterified fatty acid and 134 ± 9 μM of cholesterol. In the fasted epithelia, there was little BODIPY-staining (Fig. S6A). The entire cytosolic area of enterocytes became BODIPY-positive after perfusion for 30 min (Figs. S6E-G). Actin-staining with phalloidin indicated that there was little detectable structural damage in the epithelia with ezetimibe administration (Fig. S6B). On the other hand, alkaline phosphatase, a lipid raft marker, revealed that the treatment abolished lipid raft patches in the BBM (Figs. S6C-D). Intestinal perfusion assays in mice (Figs. 5A-B) showed that ezetimibe treatment (6–100 μg) stimulated large lipid droplet formation (Figs. 5C, S6E-P), recapitulating the results *in vitro* shown in Figs. 4, S5A-B. Tracing ^3^H-oleic acid showed that there was no difference in fatty acid uptake in the presence of ezetimibe in the small intestine (Fig. 5D). Lipid micelle perfusion in the murine intestine developed apoB accumulation in the apical perinuclear region, which was disrupted by ezetimibe (Fig. 5E). Transmission electron microscopy of jejunum sections revealed TAG-filled smooth ER and dilated Golgi membranes after 30-min luminal perfusion with ezetimibe treatment (Figs. 5F-I). For information about cellular structure, see Refs in [41, 42].

**Figure 5.**
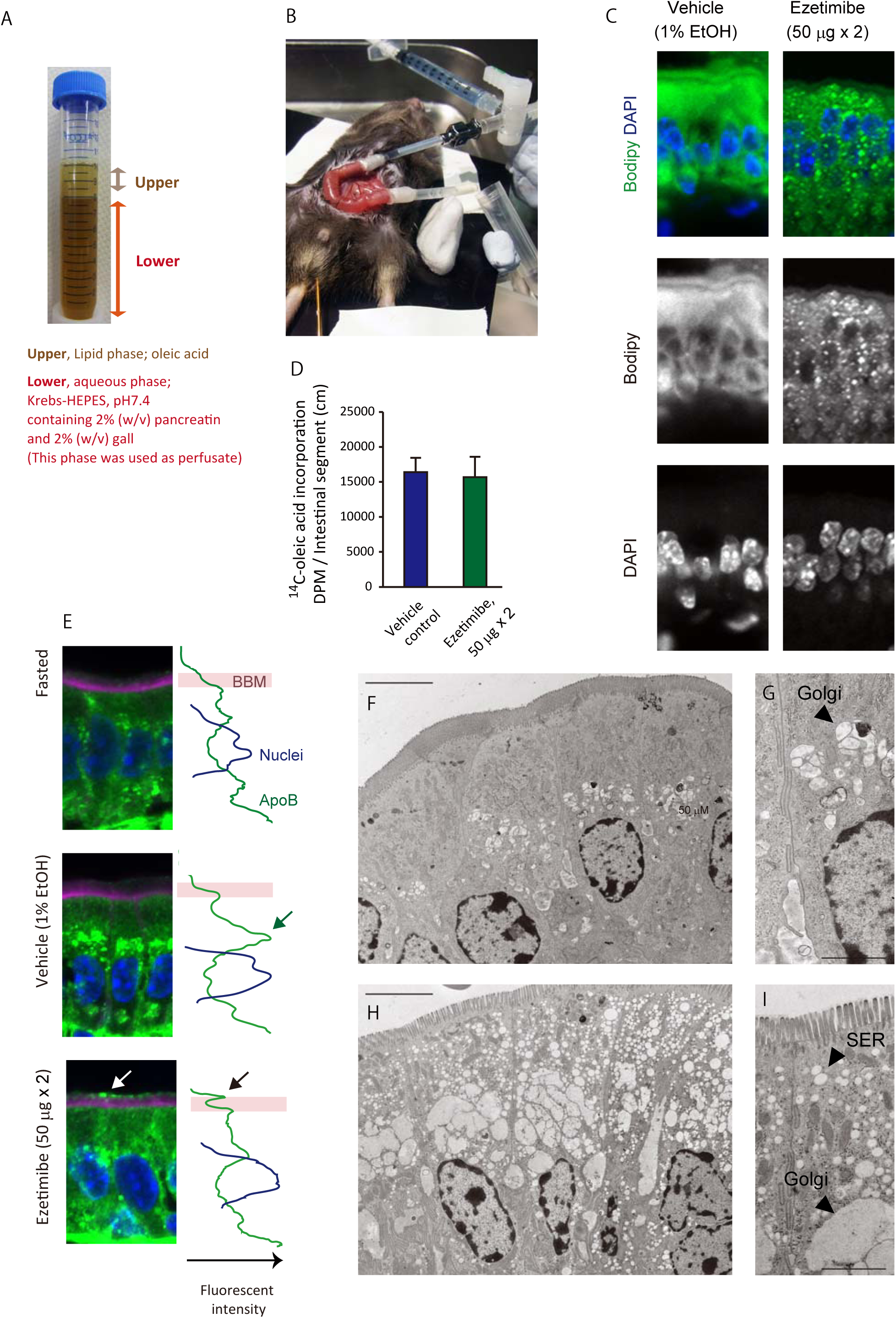
*A*, A preparation of luminal perfusate. *B* shows a mouse used for the luminal perfusion assay. *C*, ^14^C-oleic acid incorporation into the epithelia during the luminal perfusion in *B*. Total ^14^C counts were normalized by the respective lengths and are shown as mean and SEM of six replicate assays. Ezetimibe (50 μg) was given twice at from −16 to −18 h and −3 h and is as well in *D* and *E*. *D*, Ezetimibe accelerated cytosolic lipid droplet formation in the enterocytes of mice. *E*, Enterocytes accumulate apical perinuclear apoB-immunostaining with epithelial lipid load: ezetimibe disrupts the formation. Fasted, a section from a 3-h fasted mouse; vehicle, after luminal perfusion; ezetimibe, luminal perfusion after ezetimibe treatment. Lines in the side show arbitrary fluorescence intensity. *Green*, apoB immunostaining; *blue*, DAPI staining; *purple*, phalloidin for actin filaments. A green arrow indicates an accumulation of apoB in the apical perinuclear region of the enterocytes. White and black arrows indicate the presence of apoB at the tip of BBM. *F-I*, Images of murine enterocytes using transmission electron microscopy. *F* and *G*, vehicle; *H*and *I*, ezetimibe treated. Scale bars, 5 μm for *F* and *H*; 2 μm for *G* and *I*. In the presence of ezetimibe, large lipid droplets especially appeared in the apical perinuclear region in accordance with the results in Figure 4. SER, smooth endoplasmic reticulum.

### Post-prandial lipid surge was not affected by ezetimibe treatment

Ezetimibe reduced TAG-rich lipoprotein secretion *in vitro* (Fig. 1), indicating that lipid trafficking in the enterocytes was disturbed with NPC1L1 inhibition. This led us to examine whether the treatment attenuates the post-prandial lipid surge or not *in vivo*. In mice treated with a lipoprotein lipase inhibitor Tyloxapol, ezetimibe treatment did not reduce the post-prandial lipid surge significantly (Figs. 6A-B). Alternatively, we implicated that ezetimibe can alter the post-prandial lipid surge in combination with a drug that inhibits the process in the other way.

**Figure 6.**
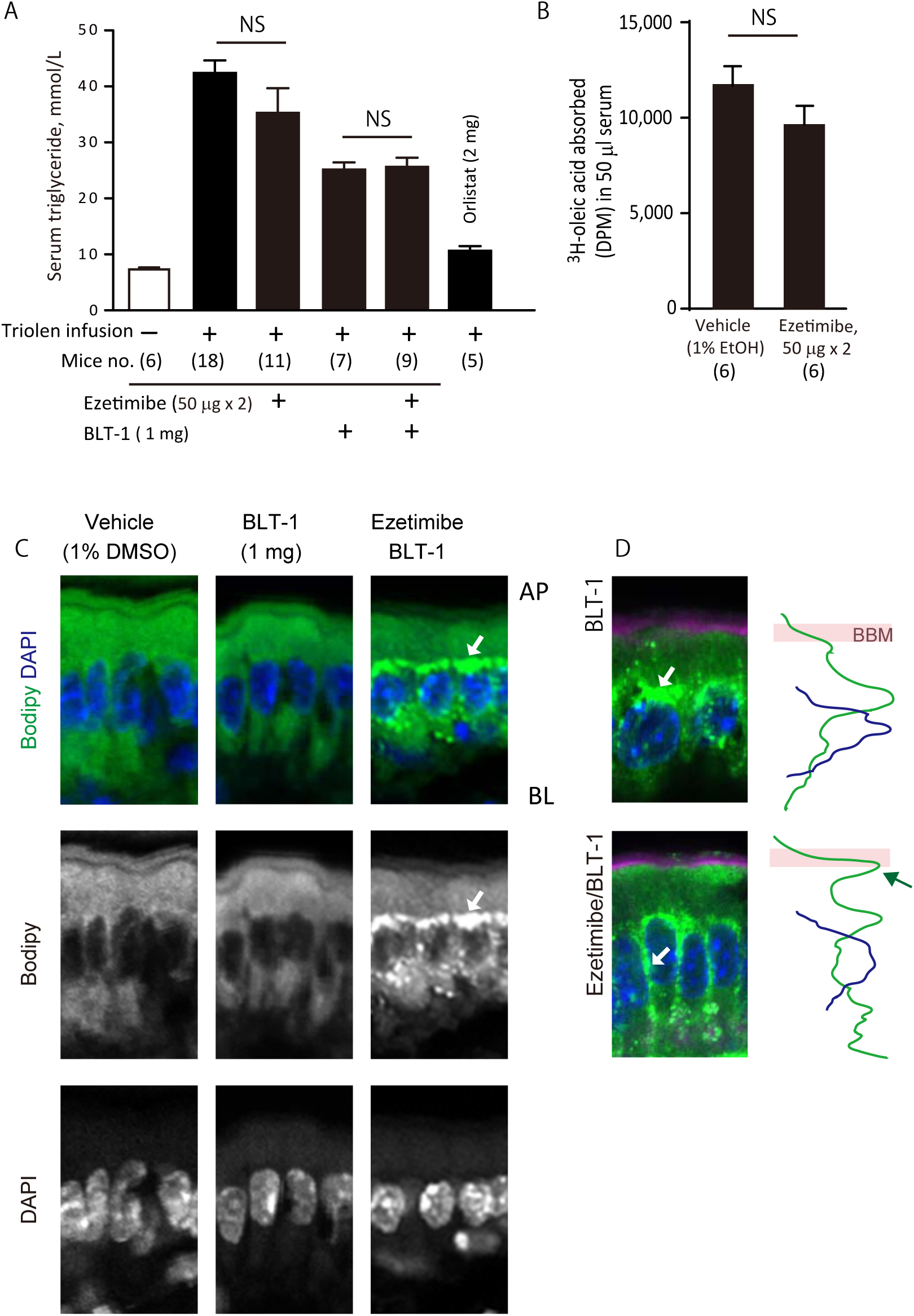
Ezetimibe did not alter triolein-induced TAG surge levels in mice. *A*, Triolein-induced TAG surge in the presence of 50 μg ×2 (at –3 h and 0 h before triolein infusion; 100 μg in total) of ezetimibe, 1 mg of BLT-1, or both. Orlistat (2 mg) was used as a positive control. NS, not significant. Data are shown mean and SEM. The number of mice used for replicates is indicated in the parentheses. *B*, ^3^H-oleic acid transition from lumen-to-circulation 3 h after the infusion. *C*, Combination of ezetimibe and BLT-1 accelerates apical perinuclear lipid droplet formation (white arrows). *D*, ApoB-immunostaining was localized around the nuclei with the combination of ezetimibe and BLT-1. Lines in the side show arbitrary fluorescent intensity. *Green*, apoB immunostaining; *blue*, DAPI staining; *purple*, phalloidin for actin filaments. The white arrow in the upper panel indicates the intense accumulation of apoB in the apical perinuclear region in the presence of BLT-1. The yellow arrow in the lower panel indicates the diffused apoB around the nuclei in the presence of ezetimibe and BLT-1. These are similar to the localization shown in Fig. 4E.

BLT-1, a scavenger receptor class B type 1 (SR-B1) inhibitor, reduces the surge after oral lipid bolus [43]. BLT-1 did not induce the formation of large lipid droplets in enterocytes in mice (Fig. 6C) and the apparent disturbance in intracellular apoB protein localization (Fig. 6D), but it reduced the post-prandial TAG surge approximately by a half (Fig. 6A), implying that BLT-1 affects the trans-intestinal lipid trafficking in a different way from ezetimibe. When mice were treated with ezetimibe in combination with BLT-1, large lipid droplets and apoB were formed around the nuclei (Figs. 6C-D), which was different from the pattern when ezetimibe was given singly (Figs. 5C-D). However, there was no synergic effect with the two compounds in the post-prandial lipid surge in mice (Fig. 6A).

## DISCUSSION

Supply of TAG and phospholipids from the lumen to enterocytes is prerequisite for chylomicron assembly and secretion. Although cholesterol is also a lipid component of the lipoprotein, the necessity in the assembly has not been explored yet. The BBM of enterocytes is rich in cholesterol and serves as its storage compartment in the cells. Moreover, enterocytes not only synthesize cholesterol *de novo* but also receive it apically (from the lumen) and basolaterally (from the blood) [17]. Thus, it has appeared to be difficult to deplete cholesterol supply in the cells experimentally, which has hampered the examination of whether or not the supply of cholesterol is dispensable for assembly.

NPC1L1 mediates cholesterol delivery from the BBM to the ER, a major cholesterol trafficking route in the enterocytes [16, 44, 45]. In the present study, we found that ezetimibe treatment activated SREBP2 (Fig. 3) and accelerated apoB degradation in Caco-2 cells (Fig. 2), likely because cholesterol shortage occurred at the ER [12, 13, 35]. In such conditions, chylomicron-like TAG-rich lipoprotein secretion by Caco-2 cells was decreased (Fig. 1), suggesting that cholesterol supply was a limiting factor for chylomicron secretory processes at least in the cells.

NPC1L1 inhibition by ezetimibe gave rise to large lipid droplets in the proximal perinuclear region in Caco-2 cells (Fig. 4) and in the murine proximal small intestine (Figs. 5, S6). These observations indicated that ezetimibe induces cholesterol depletion in murine enterocytes as well as in Caco-2 cells. Considering that SR-B1 and MTP are key for chylomicron metabolism and the inhibition results in attenuated post-prandial hyperlipidemia [46–48], we thought that BLT-1 and CP-346086, which inhibit SR-B1 and MTP, respectively, were expected to have a similar effect to ezetimibe. However, these treatments did not induce large lipid droplet formation (Figs. 6, S5). These results suggest a possibility that the lipid droplet formation is independent from lipoprotein assembly. Ezetimibe might trigger plausible checkpoints for the assembly promote lipid droplet formation alternatively. In addition, pyripyropene A, an ACAT inhibitor, and atorvastatin did not show such a lipid droplet inducible ability (Fig. S5). This rapid and easy lipid droplet formation by ezetimibe provides a unique opportunity to study the related machineries and the dynamics in the enterocytes.

Large lipid droplets emerge in enterocytes with disturbances in chylomicron assembly, secretion, or both [37–40]. Such large lipid droplets are considered to shelter absorbed alimentary TAG temporally in the cells. Accordingly, the appearance of large lipid droplets after ezetimibe treatment in mice suggested that cholesterol supply is indispensable for assembly. Furthermore, there are reports suggesting a possibility that NPC1L1 inhibition attenuates chylomicron secretion with intra-intestinal mechanisms [20, 21]. However, we did not observe any significant attenuation in the post-prandial lipid surge (Fig. 6), implying that the impact of ezetimibe is negligible in terms of attenuation of the post-prandial lipid surge. Ezetimibe reduced TAG-rich lipoprotein secretion in Caco-2 cells, but it was partial (Fig. 1). Thus, the change in lipid trafficking efficiency was not large enough to be detected by TAG measurement. Moreover, the entire small intestinal segment absorbs fatty acids. A lipid bolus we gave might not saturate the absorption capacity.

There are several limitations in this study that are worth addressing. *1)* Ezetimibe is the sole commercially available NPC1L1 inhibitor, limiting the use of a variety of chemicals to exclude possible pharmacological or chemical side effects inherent in the compound. *2)* We showed the effects of ezetimibe in enterocytes in association with lipoprotein metabolism with limited molecular identities. Cholesterol tracking, identifying associated organelles, and distribution of lipid droplet-associated molecules may be useful to understand what happens with the treatment. *3)* We measured the TAG surge in the blood, but the measurements for lymph in the thoracic duct should be sensitive for the change in epithelial lipid trafficking, but we could not perform it with technical difficulty.

Several independent research groups consistently observed that administration of ezetimibe for 2–8 weeks reduced post-prandial plasma TAG surge and apoB48 levels in volunteers who had not been treated with lipid lowering drugs [49], patients with dyslipidemia [50], hypertriglycemia [51], type-2 diabetes [52], in fructose- and fat-fed hamsters [21], and in mice fed a high-fat diet [20] (for the details, see Table S4). These may be due to extra-intestinal mechanisms, such as reduced very-low-density lipoprotein production, up-regulated clearance of remnant lipoproteins by the liver as a result of improved insulin signaling, or both [53, 54]. However, the underlying mechanisms are still obscure. Considering that, besides the liver, NPC1L1 is exclusively expressed in the small intestine, ezetimibe may act intra-intestinally to exert such favorable effects. We showed that ezetimibe altered lipid handling in the small intestinal epithelia, which may modulate lipid sensing or productivity of the signal mediators [55] to induce pleiotropic actions, such as satiety, insulin signaling modulation, intestinal motility, and so on. These elucidations are to be warranted.

In the present study, we showed that the NPC1L1 inhibitor ezetimibe disturbs lipid assimilation processes in enterocytes presumably by limiting the supply chylomicrons with cholesterol. Treatment with ezetimibe induced large lipid droplet formation in the cells, which is a unique phenomenon among tested inhibitors related to intestinal lipid metabolisms. In conclusion, NPC1L1 is likely to participate in a regulatory mechanism for the efficient chylomicron assembly and secretion by limiting intracellular cholesterol trafficking at a cellular level.

## ACKNOWLEDGEMENTS

The authors thank Dr. Sylvie Demignot (INSERM, Paris, France) for providing the Caco-2 cells and guidance regarding cell culture and lipoprotein secretion experiments, Prof. Patrick Tso (Dept. of Pathology and Laboratory Medicine, University of Cincinnati, OH, USA) for Pluronic L81, and Dr. Naonori Uozumi for critical reading of the manuscript. We are also grateful to the members of Biomedical Research Center, Saitama Medical University, for their help with the present work. This study was supported by JSPS KAKENHI Grant numbers 25504013 and 16K00864, Research Fund of Mitsukoshi Health and Welfare Foundation 2014, and Suzuken Memorial Foundation 2011, Internal Grant (No. 19-B-1-19), Saitama Medical University.

## CONFLICT OF INTEREST

The authors declare that they have no known competing financial interests or personal relationships that could have appeared to influence the work reported in this paper.

## AUTHOR CONTRIBUTIONS

**Takanari Nakano:** Conceptualization, Methodology, Investigation, Writing, Original draft preparation, Funding acquisition **Ikuo Inoue:** Conceptualization, Validation **Yasuhiro Takenaka:** Methodology **Rina Ito:** Investigation **Norihiro Kotani:** Investigation, Resources **Sato Sawako:** Investigation **Yuka Nakano:** Investigation **Masataka Hirasaki:** Investigation **Akira Shimada:** Project administration, funding acquisition **Takayuki Murakoshi:** Supervision

## Notes

### Competing Interest Statement

The authors have declared no competing interest.

