## Supplementary materials for "NPC1L1 inhibition disturbs lipid trafficking and induces large lipid droplet formation in intestinal absorptive epithelial cells"

### Supplementary Figure 1

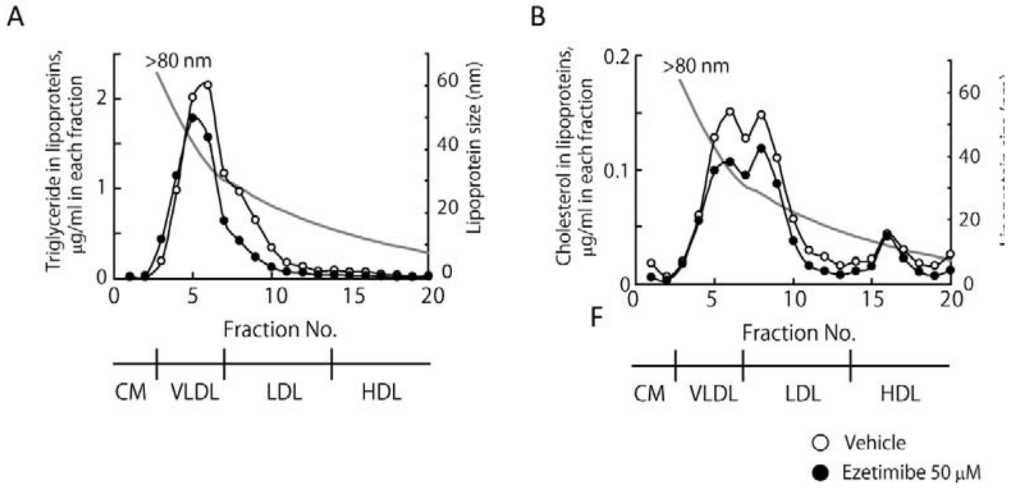


#### Lipoprotein analysis by HPLC

Lipoproteins secreted into the basolateral media were analyzed using a high-performance liquid chromatography (HPLC) column and the enzymatic detection for triglyceride and total cholesterol (Usui, S., et al. J Lipid Res 2002, 43(5): 805-814; Liposearch system, Skylight Biotech., Japan). One milliliter of the basolateral media was concentrated to 50 µl using an Amicon Ultra-4 Centrifugal Filter Unit (Merck Millipore, Billerica, MA, USA) and then applied for the HPLC analysis followed by cholesterol (*A*) and triglyceride (*B*) enzymatic detection. *Open circle*, lipid micelles only; *closed circle*, ezetimibe (50 µM). A gray line indicates lipoprotein size (right vertical axis). The estimated lipoprotein subclasses by size are shown at the bottom. Data are shown as mean of triplicate assays.

**Supplementary Figure 2**

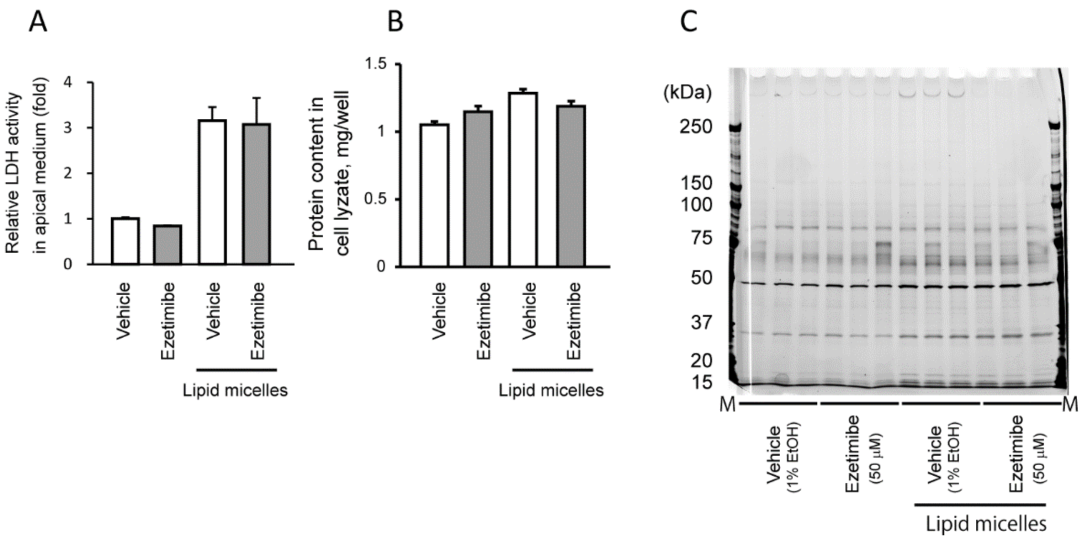


Cytotoxicity of ezetimibe in Caco-2 cells.

After 24-h incubation with or without lipid micelles, the apical and basolateral media were recovered and the cells were lyzed. *A*, Lactate dehydrogenase (LDH) activity in the apical medium (LDH cytotoxicity detection kit, TaKaRa Bio, Kusatsu, Japan). *B*, Cellular protein content. *C*, SDS-PAGE analysis of the basolateral medium followed by silver staining (2D-Silver Stain Reagent II, CosmoBio, Tokyo, Japan). Vehicle, 1% ethanol; ezetimibe, 50 μM. Data are shown as mean and SEM of triplicated assays. There was no significant difference in the band patterns between vehicle and ezetimibe treatments.

### Supplementary Figure 3

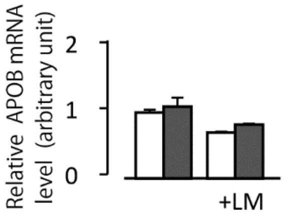


*APOB* mRNA abundance was not altered with ezetimibe in the presence or absence of lipid micelles at 6 h after ezetimibe treatment, as examined using Student's *t*-test. Data are shown as mean and SEM of four replicate assays. +LM, lipid micelles added.

**Supplementary Figure 4**

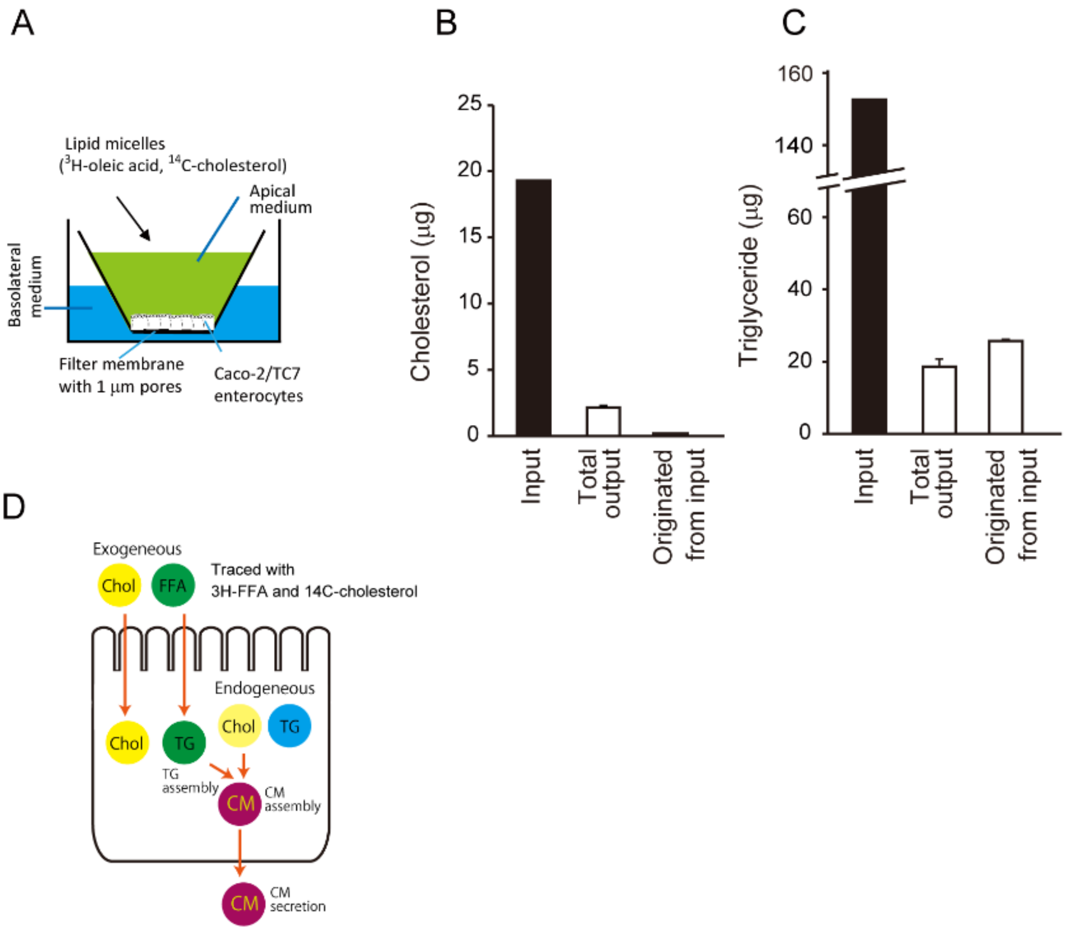


Differentiated Caco-2 cells secrete triglyceride-rich lipoproteins predominantly containing endogenous cholesterol, but not that taken up apically.

*A,* A schematic presentation of the culture system for Caco-2 cells. Lipid tracers were given via the apical medium (*green*). The basolateral medium (*blue*) was analyzed for secreted lipoprotein. *B and C*, Cholesterol supplied apically is not appeared in triglyceride-rich lipoproteins in differentiated Caco2 cells (B). On the other hand, $^3$H-oleic acid measurement shows that triglyceride in secreted lipoproteins originates from the apically supplied fatty acids. Cholesterol (*left*) and oleic acid (as triglyceride in the basolateral media) (*right*) given apically were traced quantitatively (input and total output) or by tracer activity (originating from the input). *Total output*, amounts of total secreted lipid from Caco-2 cells determined in quantitative assays; *Originated from input*, amounts of secreted lipids originating from added lipid micelles calculated by movement of cholesterol and oleic acid tracers. *D*, Schematic presentation of lipid origins for chylomicron assembly in enterocytes.

### Supplementary Figure 5

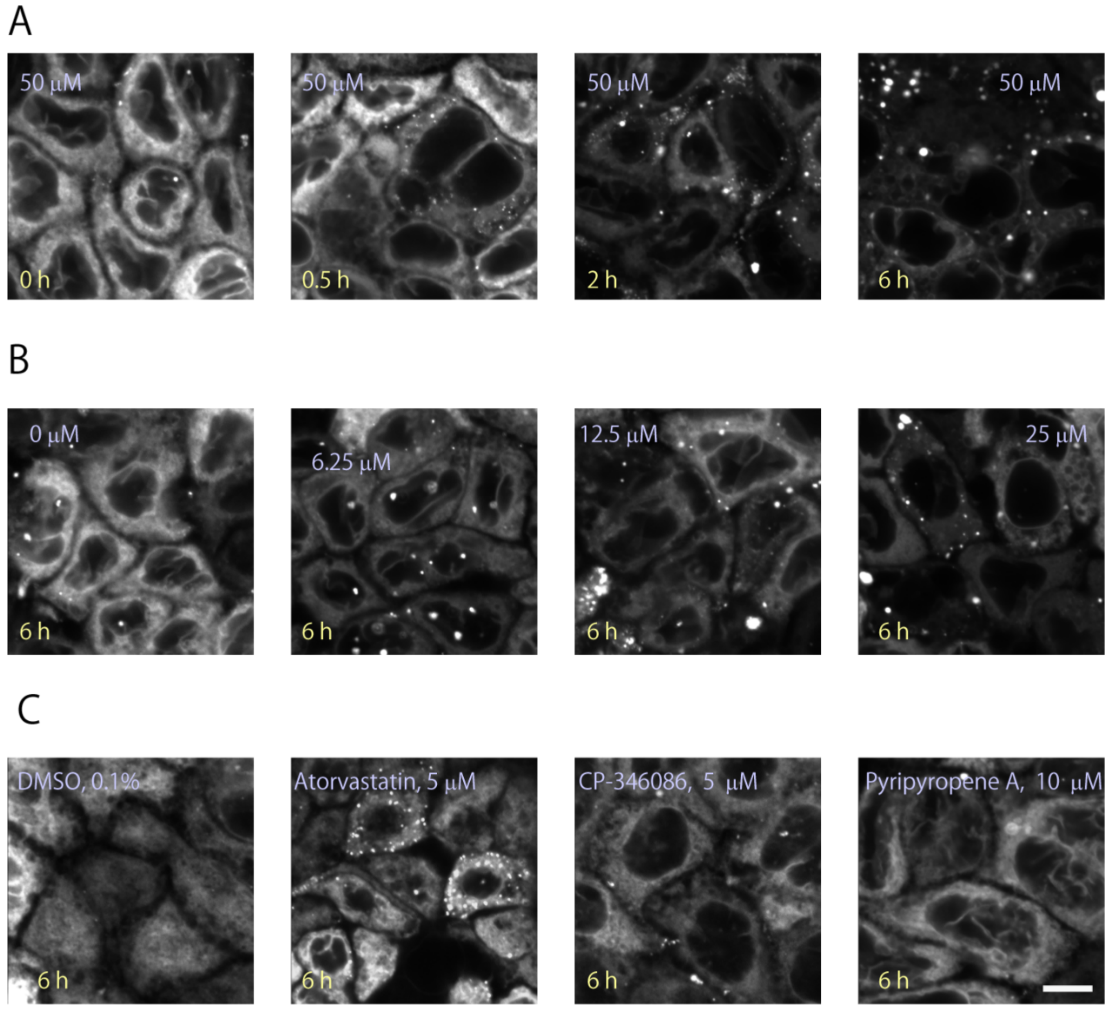


*A,* Effect of duration of ezetimibe treatment (0.5–6 h) on lipid droplet formation. Ezetimibe (50 µM) was applied, and BODIPY™ 493/503 staining was examined at the indicated time points. Dispersed BODIPY staining in the cytosolic area gradually disappeared with time, and large lipid droplets formed.

*B*, Dose-dependent effect of ezetimibe treatment on lipid droplet formation. Apparent large lipid droplet formation was observed at doses greater than 12.5 µM.

*C*, Three other lipid metabolism inhibitors did not show effects comparable to ezetimibe. Atorvastatin, HMG-CoA dehydrogenase inhibitor; CP-346086, microsomal triglyceride transfer protein inhibitor; pyripyropene A, acyl-CoA acyltransferase inhibitor.

### Supplementary Figure 6

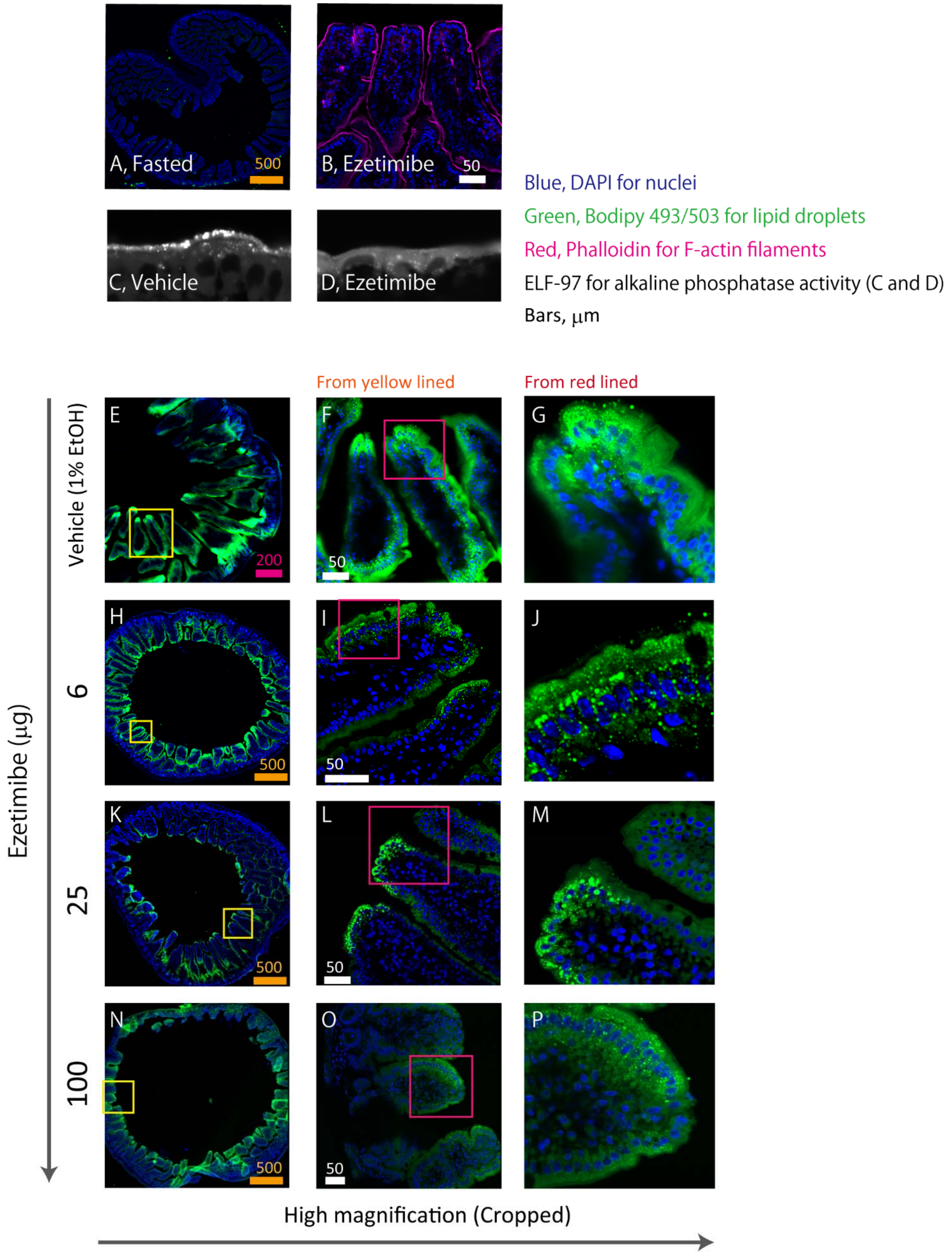

**Supplementary Figure 6 (*continued*)**

Ezetimibe stimulates large lipid droplet formation in enterocytes in mice.

*A*, The jejunum obtained from mice fasted overnight showed no BODIPY staining.

*B*, Ezetimibe treatment (100 μg) did not affect microvillus appearance, as examined using F-actin staining.

*C* and *D,* Alkaline phosphatase activity staining using ELF-97 indicated the presence of lipid raft patches at the brush border membrane, and these lipid rafts disappeared with ezetimibe treatment (*D*).

*E-P*, BODIPY 493/503 staining for the jejunal segments after luminal perfusion assay. As low as 6 μg ezetimibe (3 μg ×2) was sufficient to stimulate lipid droplet formation. The yellow or red rectangles indicates the areas cropped. Ezetimibe was given twice. The amounts of ezetimibe are shown as totals.

### Table S1. Chemicals used in this study

| Reagent name | Provider | Identifier |
|---|---|---|
| Alexa Fluor 568 phalloidin | Molecular Probes | A12380 |
| Atorvastatin | Tocris Biosciences | 3776 |
| BODIPY 493/503 | Molecular Probes | D3922 |
| Cholesterol | Nacalai Tesque | 08721-62 |
| Oleic acid | Nacalai Tesque | 25630-51 |
| Colhicine | Nacalai Tesque | 09305-05 |
| CP-346086 | Sigma-Aldrich | PZ0103 |
| DAPI | Nacalai Tesque | 11034-56 |
| Ezetimibe | Santa Cruz | sc-205690 |
| Gall | FUJIFILM Wako Pure Chemical Corp. | 8008-63-7 |
| L-α-Lysophosphatidylcholine, from Egg Yolk | FUJIFILM Wako Pure Chemical Corp. | 123-03781 |
| Monencin | Tocris Biosciences | 5223/500 |
| Nocodazole | Abcam | ab120630 |
| Pancreatin | Nacalai Tesque | 25930-34 |
| Protease inhibitor cocktail | Sigma-Aldrich | P8340 |
| Pyripyropene A | Biolinks | BLK-0570 |
| Taurocholate | MP Biomedicals | 102930 |
| Triolein | Nacalai Tesque | 35227-12 |
| Tunicamycin | Santa Cruz | sc-3506 |

**Table S2. Antibodies used in this study**

| Category | Immunogen | Reactivity | Host | Provider | Identifier | Labeling | Dilution | Figure |
|---|---|---|---|---|---|---|---|---|
| Western blotting | Apolipoprotein B | Human | Rabbit | The Binding Site | PP086 | HRP | 1:1,000 | 1D |
| | SREBP1 | Human | Rabbit | Santa Cruz | C-20 | | 1:400 | 3C |
| | SREBP2 | Human | Rabbit | Abcam | ab30682 | | 1:200 | 3C |
| | β-actin | | Mouse | Abcam | ab6276 | | 1:5,000 | 3C |
| | IgG | Rabbit | Goat | GE Healthcare | N934 | HRP | 1:1,000 | 1D/3C |
| | IgG | Mouse | Rabbit | Sigma-Aldrich | A9044 | HRP | 1:1,000 | 3C |
| Immunoprecipitation | Apolipoprotein B | Human | Rabbit | Biodesign International Inc. | K45253G | | 1:50 | 2B/2C |
| | Apolipoprotein B | Human | Rabbit | Chemicon | AB742 | | 1:50 | 2B |
| | Apolipoprotein B | Human | Rabbit | The Binding Site | PC086 | | 1:50 | 2B |
| Immunohistochemistry | Apolipoprotein B | Mouse | Rabbit | Merdian Life Science Inc. | K23300R | | 1:500 | 5E/6D |
| | IgG | Rabbit | Goat | ThermoFisher Scientific | A11070 | AlexaFlour-488 | 1:500 | 5E/6D |

**Table S3. Primer pairs used in this study.**

| HGNC symbol | Accession No. | Name | Forward primer (5'-3') | Reverse primer (5'-3') | Refs. |
|---|---|---|---|---|---|
| *ABCA1* | NM_005502.2 | ATP-binding cassette A1 | ATGTCCAGTCCAGTAATGGTTCTGT | CGAGATATGGTCCGGATTGC | (1) |
| *ACACB* | NM_001093.3 | Acetyl-CoA carboxylase β | CAGAGCATCGTGCAGTTGGT | TGCTCAACACGCAAGTATCTTCTC | (2) |
| *ACAT2* | NM_005891.2 | Acetyl-CoA acetyltransferase 2 | GAGACTTACCCTAGGACGCCCT | AGTTCTTGGCCACATAATTCCAC | (3) |
| *APOB* | NM_000384 | Apolipoprotein B (including Ag(x) antigen) | ACCTCCAGAACATGGGATTGC | GGGCTGGTGTCCTAACAGTC | (4) |
| *FAS* | NM_004104.4 | Fatty acid synthase | TATGCTTCTTCGTGCAGCAGTT | GCTGCCACACGCTCCTCTAG | (5) |
| *GPAT1* | NM_006411.3 | 1-acylglycerol-3-phosphate *O*-acyltransferase 1 | GGAAAGTTTATCCAGTATGGCATT | TGATATCTTCCTGGTCATCGTG | *1 |
| *HPRT1* | NM_000194 | Hypoxanthine phosphoribosyltransferase I | GTAATTGGTGGAGATGATCTCTCAACT | TGTTTTGCCAGTGTCAATTATATCTTC | (6) |
| *INSIG1* | NM_005542.4 | Insulin induced gene 1 | ATCTTTTCCTCCGCCTGGT | GGGGTACAGTAGGCCAACAA | *1 |
| *LDLR* | NM_000527.4 | Low-density lipoprotein receptor | CAGATATCATCAACGAAGC | CCTCTCACACCAGTTCACTCC | (7) |
| *MTTP* | NM_000253.2 | Microsomal triglyceride transfer protein | AGCACCTCAGGACTGCGAAGA | CAGAGGTGACAGCATCCACCA | (8) |
| *MVD* | NM_002461.1 | Mevalonate (diphospho) decarboxylase | AACATCGCGGTCATCAAGTA | TTAACTGGTCCTGGTGCAGAG | *1 |
| *NPC1L1* | NM_013389.2 | NPC1 (Niemann-Pick disease, type C1, gene)-like 1 | ACATCAGCGTGGGACTGG | AGTCAAGCAGGTACGAGTCCTT | *1 |
| *PMVK* | NM_006556.3 | Phosphomevalonate kinase | TTTTGCAGGAAGATTGTGGA | CTCCGTGTGTCACTCACCA | *1 |
| *SCD1* | NM_005063.4 | Stearoyl-Coenzyme A desaturase 1 | CCTACCTGCAAGTTCTACACCTG | GACGATGAGCTCCTGCTGTT | *1 |

*1, The primer sets were designed with using the Assay Design Center at http://www.roche-applied-science.com/.

References; the sources for primer sequences in Table S2

Table S4. Studies examining the effect of ezetimibe on post-prandial triacylglycerol (TAG) increase

| Studies | Masuda D (2009) | Yunoki K (2011) | Bozzetto L (2011) | Hiramitsu S (2012) | Naples M (2012) | Kikuchi (2012) | Sandoval JC (2010) | |
|---|---|---|---|---|---|---|---|---|
| Subjects | 10 | 20 | 15 | 20 | Hamster | 20 | Mice (WT) | Mice (CD36KO) |
| Subject conditions | Type IIb hyperlipidemia | Volunteers | Type-2 diabetes | Obesity or hypertriglycaemia | FF diet | Obese/dyslipidaemia | Western diet (WT) | Western diet (CD36KO) |
| Evaluation | Post-prandial AUC-TG change | Post-prandial AUC-TG change | Chylomicron TG at 6 h | Post-prandial AUC-TG change | TG(mM) AUC for 3 h | | Lipid transit by tracer at 3 h after the infusion (lymph duct) | |
| Study settings | Randomized | Randomized | Double-blind cross-over design | Before-after study | In experimental animals | Randomized/crossover | In experimental animals | |
| Administration | 10 mg per day for 2 months | 10 mg per day for 4 wks | 10 mg per day for 6 wks (simvastatin 20mg per day for both groups) | 10 mg per day for 4 wks | 1-2 mg/kg per day for 2wks (in chow) | 10 mg per day for 4 wks | 10 mg/kg per day for 3 wks (in chow) | |
| Results | | | | | | | | |
| Reference value | 2448 | 1419 | 29 | 1950 | 36.3 | | 154000 | 46800 |
| Ezetimibe treated | 1863 | 968 | 18 | 1523 | 23.1 | | 56300 | 34800 |
| Order | AUC TG | mg h/dl | mg/dl x 6 h AUC | AUC TG | AUC TRL-TG | AUC TG (mg/dl) | dpm/ml ($^3$H-triolein) | dpm/ml ($^3$H-triolein) |
| Reduction (%) | 24% | 32% | 38% | 18% | 37% | 27% | 63% | 26% |

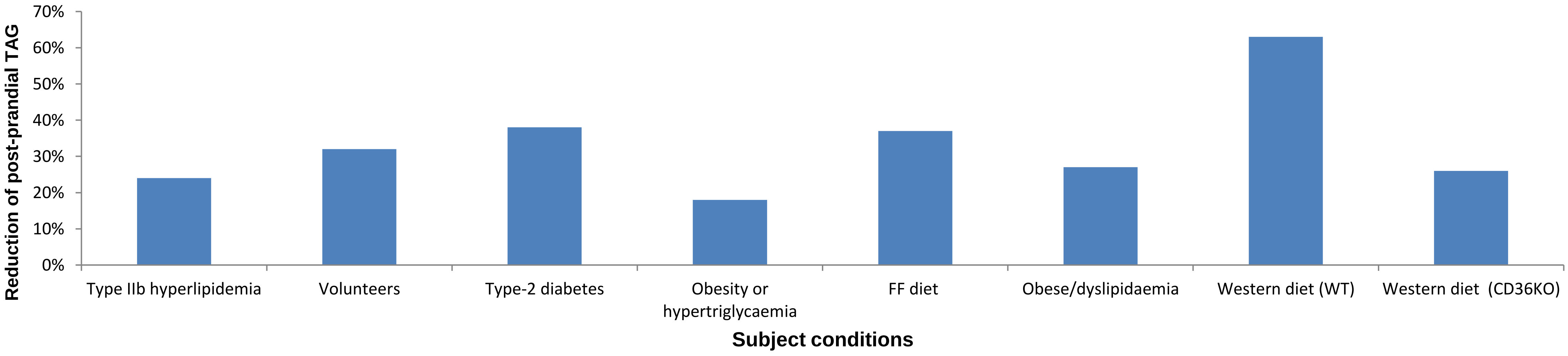


**Abbreviations**: AUC, area under the curve; CD36KO, CD36 knockout mice; dpm, decay per minute; FF, fructose and fat; h, hour; TAG, triacylglycerol; wks, weeks; TRL, triacylglycerol-rich lipoproteins; WT, wild-type
